# Social Anxiety Alters Theory of Mind Activation and Intersubject Neural Variability During Movie Viewing

**DOI:** 10.1101/2024.11.12.622970

**Authors:** Margot Mangnus, Saskia B. J. Koch, Robin Devillers, Peter Hagoort, Jana Bašnáková, Arjen Stolk

## Abstract

Social anxiety is characterized by an intense fear of judgment in social situations, yet the underlying mechanisms driving this condition remain poorly understood. One hypothesis holds that specific alterations in Theory of Mind (ToM) affect the ability to interpret others’ thoughts and emotions. Another hypothesis proposes that broader interpretive biases lead individuals to perceive social cues as overly significant, even in neutral settings. We investigated these possibilities by measuring brain activity, pupil responses, and heart rates in socially anxious individuals and matched controls as they viewed ‘Partly Cloudy’, an animated film known to engage the ToM network during specific scenes. While overall brain activity during ToM-related scenes was similar across groups, socially anxious participants exhibited reduced activation in the left posterior superior temporal sulcus (pSTS), a key area for ToM processing. Additionally, intersubject correlation analysis revealed a distinct neural response pattern in the socially anxious group, marked by uniform responses in sensory regions and heightened variability in higher-order cortical areas. This pattern persisted throughout the film and occurred without changes in heart rate or pupil responses, indicating a neural processing bias that manifests even in non-evaluative settings. These findings provide a neural basis for ToM alterations and broader interpretive biases in social anxiety, supporting cognitive-behavioral models and suggesting novel targets for intervention.

## Introduction

Social Anxiety Disorder (SAD), also known as social phobia, is defined by an intense and persistent fear of social situations, often manifesting during childhood or adolescence. Affecting approximately 12% of adults at some point in their lives [1], SAD leads individuals to experience excessive worry and to avoid situations where they might be judged or scrutinized, from everyday interactions to high-stakes events like public speaking or job interviews [2]. While avoidance might seem protective, it typically intensifies feelings of isolation and deteriorates overall mental health [3], highlighting the urgent need for a deeper understanding of the mechanisms behind this anxiety disorder to develop more effective treatments.

This study investigates two hypotheses concerning the underlying mechanisms of social anxiety. The first hypothesis posits that alterations in Theory of Mind (ToM) affect individuals’ ability to accurately interpret others’ thoughts and emotions [4], thereby increasing stress and anxiety during social interactions [5, 6]. Evidence indicates that individuals with SAD typically underperform on ToM tasks, such as the Reading the Mind in the Eyes Test and the Movie Assessment of Social Cognition, relative to nonclinical controls [7–10]. Furthermore, research suggests that individuals with SAD may perceive emotions as more intense than they actually are, raising questions about the potential over-or under-utilization of their ToM capabilities [4, 11, 12]. Despite these findings, neuroimaging studies exploring ToM in the context of social anxiety are limited [13]. It also remains unclear whether alterations in ToM are a constant feature of the disorder or are predominantly triggered by situations imposing explicit task demands and evaluative pressures. Clarifying this distinction could help determine whether these ToM changes are inherent to SAD or contextually induced.

The second hypothesis suggests that individuals with SAD possess broader interpretive biases, perceiving social cues as overly significant with profound personal implications [14, 15]. While theoretical models vary, they generally agree that these biases compel individuals to incessantly monitor their environments for potential social evaluative threats [16–20] and to critically assess their own behaviors [21, 22]. Neuroscientific studies support these theories by demonstrating that feedback from social performance tasks, such as public speaking, elicits negatively biased responses in brain areas like the precuneus and frontoparietal regions, reinforcing a negative self-image [23]. Additionally, research on children with social anxiety shows considerable intersubject variability in these regions when exposed to socioemotionally charged films, suggesting a heightened subjective response to social stimuli even in task-free settings [24]. Nonetheless, it remains uncertain whether these brain responses are linked to ToM processes or reflect distinct interpretive mechanisms. Although activation in these areas hints at potential overlap with the ToM network, which includes the medial prefrontal cortex, precuneus, temporoparietal junction, and posterior superior temporal sulcus, direct comparisons of these hypotheses through functional brain imaging have yet to be conducted.

In this study, we employed Pixar’s animated short film ‘Partly Cloudy’ to investigate how individuals with varying levels of social anxiety utilize their ToM capabilities in a task-free environment. The film, which portrays the developing friendship and interactions between a stork and a cloud, contains specific scenes known to activate the ToM network by illustrating characters’ mental states [25–27]. Beyond analyzing responses to these ToM-specific scenes, the film’s continuously evolving narrative provides an effective platform for assessing individual differences in processing identical stimuli throughout the film using dynamic intersubject correlation analysis. Previous research using this method found reduced intersubject variability in brain and pupil responses among autistic individuals during substantial portions of the film [28]. This uniformity was primarily observed in brain regions outside the ToM network, while the ToM network itself exhibited robust responses to scenes depicting mental states in both autistic and non-autistic viewers. These observations highlight the film’s utility in examining both ToM-specific activation and broader interpretive processing. Leveraging these insights, the current neuroimaging study aims to delineate how these processes interact and manifest in individuals with social anxiety under comparable viewing conditions.

Forty-three socially anxious participants and 43 matched controls viewed the film inside an MRI scanner. To minimize performance-related anxiety, we measured whole-brain activity and pupil responses without providing specific task instructions, and we monitored heart rate to assess physiological arousal levels [29]. After the viewing, participants completed an unannounced questionnaire evaluating their engagement with the film. Our analysis comprised two primary components. First, we conducted event-related comparisons between groups during scenes that depicted characters’ mental states, assessing both functional and anatomical aspects of ToM activation in individuals with high and low levels of social anxiety. Based on previous observations of diminished ToM performance [8], we anticipated altered activation within the ToM network among socially anxious individuals during these scenes. Second, we analyzed intersubject correlations throughout the film, focusing on sustained variations that could indicate differences in subjective engagement or interpretive biases among socially anxious individuals. Drawing on related research involving children [24], we hypothesized that socially anxious participants would show increased intersubject variability in frontoparietal regions associated with socioemotional processing. Because results indicated heightened neural variability in regions previously showing reduced variability in autism, we compared neural patterns between these groups to uncover common and distinct mechanisms underlying social processing in autism and social anxiety [30].

## Materials and methods

### Participants

Eighty-six adult participants were recruited through Radboud University’s database, social media advertisements, and campus postings. Exclusion criteria included the use of psychotropic or systemic glucocorticoid medications, systemic diseases, severe cognitive impairments, or ongoing neurological treatments. Control participants additionally had no current psychiatric diagnoses. Participants were matched for gender, age, and both verbal and nonverbal IQ scores [31, 32] (see Table 1). Social anxiety levels were assessed using the Liebowitz Social Anxiety Scale (LSAS) [33, 34], a 24-item self-report instrument where participants rated their fear and avoidance behaviors on a Likert scale from zero (none/never) to three (severe/usually) across anxiety-inducing situations (e.g., eating in public, interacting with strangers). A cutoff score of 30 [35] categorized participants into high social anxiety (LSAS ≥ 30) and control (LSAS < 30) groups, each comprising 43 individuals. Data collection included MRI scans (*n* = 86), pupillometry (*n* = 75), and heart rate measurements (*n* = 74) while participants viewed the film, followed by a post-viewing questionnaire (*n* = 84). All participants provided written informed consent in accordance with local ethics guidelines (CMO region Arnhem-Nijmegen, the Netherlands, file number 2019-6059) and received compensation for participation.

**Table 1.** Demographic data.

|  | *SA Group* | *CON Group* | *Group Difference* |
| --- | --- | --- | --- |
| N (male:female) | 43 (16:27) | 43 (18:25) | $\chi^2 (1, N = 86) = -0.05, p = .83$ |
| Age (years) | 26.3 (65.9) | 26.0 (5.2) | $t (84) = 0.23, p = .82$ |
| Verbal IQ (WAIS-III) | 124 (14) | 128 (16) | $t (84) = -1.36, p = .17$ |
| Nonverbal IQ (RPM) | 102 (11) | 102 (11) | $t (84) = -0.23, p = .82$ |
| Social Anxiety (LSAS) | 51.4 (18.4) | 15.2 (7.5) | $t (84) = 11.8, p < .001$ |

Values are given as Mean (Standard Deviation). SA, Social Anxiety; CON, Control.

### Experimental protocol

Participants watched the nonverbal animated film Partly Cloudy (5 minutes 45 seconds), depicting the developing friendship between a stork and a cloud. The film was annotated for three event types: Mental (44 seconds across 4 scenes), Pain (26 seconds over 7 scenes), and Control (24 seconds across 3 scenes), as shown in Fig. 1 [25]. Mental events were expected to elicit Theory of Mind (ToM) inferences about characters’ thoughts and emotions, such as distress or betrayal. Pain events depicted physical discomfort, like electric shocks or animal attacks. Control events showed neutral content, such as birds in flight or cloudscapes without foreground characters. These categories were used to analyze ToM-related brain and pupil responses. Before the MRI session, participants were acclimated to the MRI environment using a dummy scanner to reduce anxiety. After viewing the film, they completed an undisclosed questionnaire describing the plot in their own words. Independent raters analyzed these summaries using a taxonomy of mental state terms [36], categorizing words referencing mental states and emotions versus general content. An independent *t*-test compared the frequency of mental state-related words between groups. To test for a potential negativity bias among socially anxious participants, a quasi-Poisson regression model assessed the ratio of negative emotion words to the total number of emotion-related words used by each participant.

**Fig. 1.**
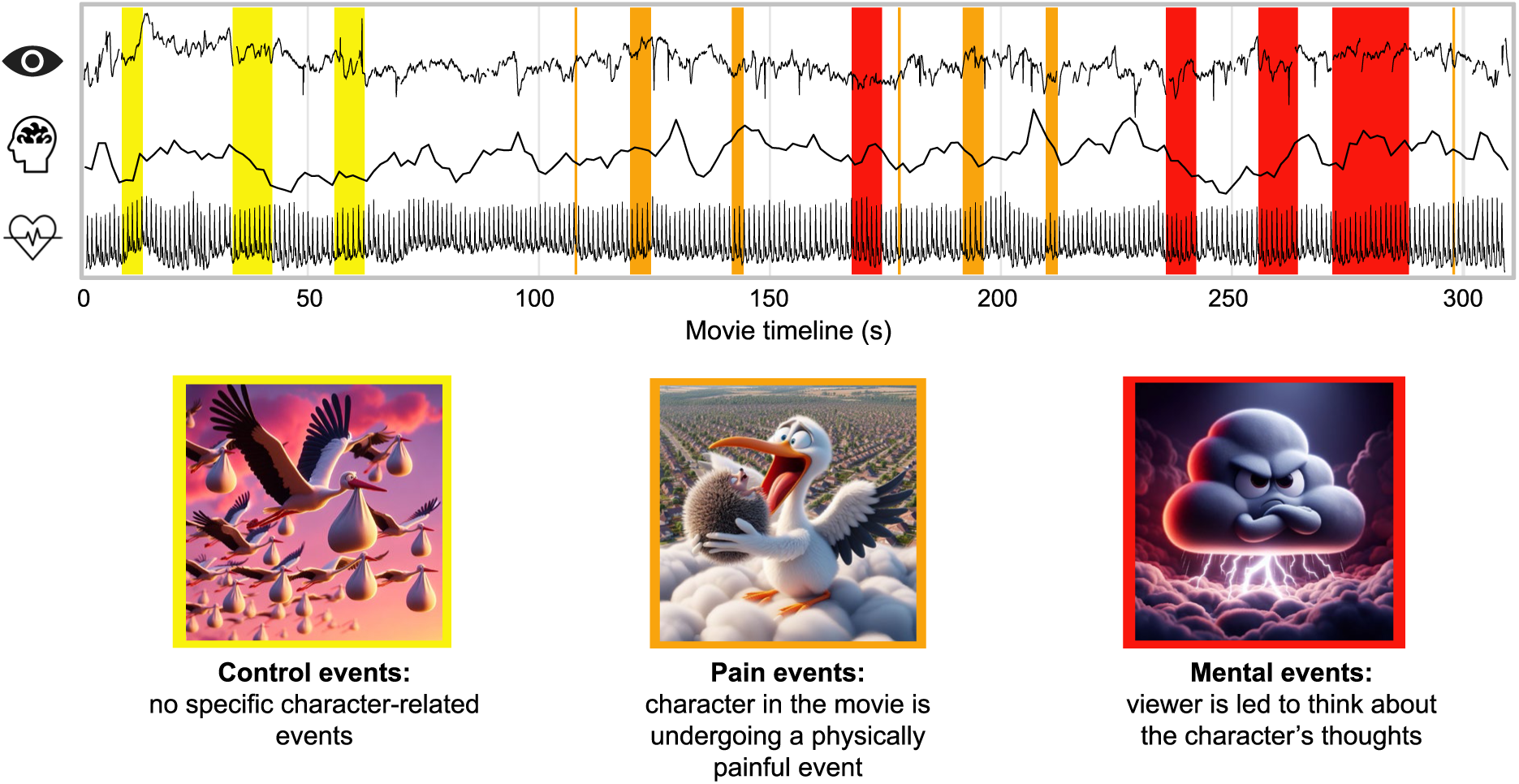
Experimental stimuli and responses. Socially anxious participants and matched controls viewed the animated short ‘Partly Cloudy,’ which portrays the developing friendship and interactions between a stork and a cloud. During the viewing, participants’ pupil responses, brain activity, and heart rates were continuously recorded. The film is annotated to highlight three distinct event types: Mental, Pain and Control, each marked on the film’s timeline. Mental events are expected to engage viewers’ Theory of Mind, prompting them to infer the characters’ thoughts and emotions. Pain events illustrate the characters experiencing physical discomfort, while Control events feature only passive or background imagery. The images displayed were generated using Bing’s Copilot to closely replicate scenes from the film, adhering to copyright constraints.

### Data acquisition

During the film screening, heart rate was continuously recorded at 5000 Hz using a finger pulse sensor connected to a BrainAmp ExG MR amplifier (BrainVision software), and pupil diameter was monitored at 1000 Hz using an Eyelink 1000 Plus eye-tracker. MRI scans were conducted on a Siemens 3T scanner with a 32-channel head coil. High-resolution structural images were obtained using a T1 MPRAGE sequence (TR = 2200 ms, TI = 1100 ms, TE = 2.6 ms, flip angle = 11°, voxel size = 0.8 mm isotropic, and acceleration factor = 2). Functional images were acquired using a multi-band multi-echo sequence (TR = 1500 ms, TEs = 13.4/34.8/56.2 ms, flip angle = 75°, voxel size = 2.5 mm isotropic, and acceleration factor = 2). Motion analysis revealed no significant differences between socially anxious and control participants in framewise displacement (mean FD = 0.16 ± 0.08 vs. 0.16 ± 0.07, *t*(84) = 0.20, *p* = .84; max FD = 0.09 ± 1.28 vs. 1.15 ± 1.13, *t*(84) = 0.67, *p* = .51) or total head motion, calculated from translation and rotation parameters during realignment (translation: 118.4 ± 110.4 vs. 107.4 ± 78.7, *t*(84) = –0.53, *p* = .59; rotation: 2.3 ± 2.0 vs. 2.4 ± 1.9, *t*(84) = 0.18, *p* = .86).

### Heart rate analysis

Heart rate data were preprocessed by removing scanner-induced artifacts using a deconvolution filter from the BrainAmpConverter toolbox. The cleaned signal was then band-pass filtered between 0.2 Hz and 3 Hz to retain biologically relevant frequencies [37]. Heartbeat peaks were detected within 600 ms intervals, and heart rate metrics were derived from the intervals between these peaks. Intersubject variability in heart rate responses was assessed using the dynamic intersubject correlation method described in the pupillometry analysis section.

### Pupillometry analysis

Pupillometry data were preprocessed to extract changes in pupil size, an indicator of cognitive effort [38]. Blinks and saccades were removed using noise-based detection and adaptive velocity threshold algorithms [39, 40], while visual inspections eliminated squints and gaze jumps [41]. Each participant’s pupil timeseries was standardized (*z*-scored) and adjusted for global luminance variations using a 5th-order polynomial model in R [42], with luminance derived from RGB values per the Rec. 709 standard [43]. Group differences in mean pupil responses across event types were assessed using a 3×2 mixed-design ANOVA, with event type (*Mental*, *Pain*, *Control*) as the within-subject factor and group (socially anxious, control) as the between-subject factor. Significant interaction effects (*p* < .05) were analyzed using Tukey’s Honest Significant Difference tests.

To examine intersubject variability, we conducted dynamic intersubject correlation analysis on the pupil timeseries. Using a leave-one-out method [44], each participant’s timeseries was correlated with the average of all other group members, applying a 30-second sliding window with 100 ms steps to generate continuous correlation timeseries spanning the entire movie. The correlations were Fisher *z*-transformed and subjected to a nonparametric cluster-based permutation test to evaluate differences in pupil response variability between groups [45]. This test addresses the multiple comparisons problem in timeseries analysis by aggregating significant adjacent data points into clusters. These clusters are tested against a null distribution formed by randomly shuffling participant labels and reculating statistics, effectively controlling for false positives. Statistical significance was assessed using a two-sided independent samples *t*-test, with 10,000 permutations. Clusters with a Monte-Carlo *p*-value of .05 or less were considered statistically significant.

### fMRI analysis

fMRI data were preprocessed and analyzed using SPM12 (https://www.fil.ion.ucl.ac.uk/spm). Echoes were merged into a single volume, and images were realigned to the first image using 2nd degree B-spline interpolation with six rigid-body transformation parameters. Spatial distortions were minimized by unwarping images using a fieldmap acquired alongside the functional images. The mean functional images were coregistered with anatomical images and normalized to MNI space using tissue probability maps. Images were then spatially smoothed with an 8 mm full-width at half-maximum kernel. The first-level analysis included regressors for Mental, Pain, and Control events, end credits, head movement parameters (including squared, cubic terms, and derivatives), and signals from white matter, cerebrospinal fluid, and out-of-brain voxels.

To assess group differences in ToM activation, two separate 2×2 mixed-design ANOVAs were conducted. *Mental* and *Pain* events or *Mental* and *Control* events served as within-subject factors, and participant group (*socially anxious*, *control*) was the between-subject factor. We hypothesized that these contrasts would reveal common activations within the ToM network, isolated using conjunction analysis and specifically masked to emphasize ToM activation in the control group. Results are presented as whole-brain cluster-level corrected effects (*p_FWE_* < .05, initial cluster-forming threshold at *p* < .001), with anatomical locations identified using the SPM Anatomy Toolbox [46]. A-region of-interest (ROI) analysis focused on key ToM-related brain regions, including the right temporoparietal junction (rTPJ), precuneus, and medial prefrontal cortex (mPFC; Schurz et al., 2014), defined by 8 mm radius spheres at peak voxels from the combined ToM contrasts. Bayesian analysis assessed variations in ToM network activation between groups by calculating Bayes Factors (BFs) to determine the strength of evidence for the null hypothesis versus alternative models [48].

To examine differences in intersubject variability at the neural level, we performed dynamic intersubject correlation analysis on voxel timeseries, adjusted for head movement and tissue signals. Using a combined leave-one-out and sliding-window approach similar to that used in the heart rate and pupillometry analyses, we generated voxel correlation timeseries. For computational efficiency, the data were spatially and temporally downsampled by a factor of three, resulting in a voxel resolution of 7.5 mm and a sampling interval of 4.5 seconds. These data underwent a nonparametric cluster-based permutation test [28], correcting for multiple comparisons across voxels and timepoints. To visualize the spatial distributions of identified spatiotemporal clusters, we aggregated the *t*-values across all timepoints, producing a three-dimensional representation highlighting areas with significant variability differences throughout the film’s duration. Lastly, we quantified the overlap between the correlation clusters and the ToM network by assessing their spatial overlap relative to the total voxel size of the detected clusters.

## Results

### Post-viewing movie assessment

After viewing the movie, both socially anxious and matched control participants used a similar frequency of words related to mental states (M_SA_ = 0.053, M_CON_ = 0.048, *t*(82) = 0.47, *p* = .64; Fig. 2a and b) and emotions (M_SA_ = 0.042, M_CON_ = 0.034, *t*(82) = 0.74, *p* = .46) when describing the plot. However, socially anxious participants used a significantly lower proportion of negatively charged emotion words (M_SA_ = 0.65, M_CON_ = 0.85, *t*(50) = –2.61, *p* = .012; Fig. 2c). These results suggest that, although both groups similarly recognized the mental states depicted in the film, socially anxious individuals may have adopted a more positive perspective on the plot upon reflection than their non-anxious counterparts.

**Fig. 2.**
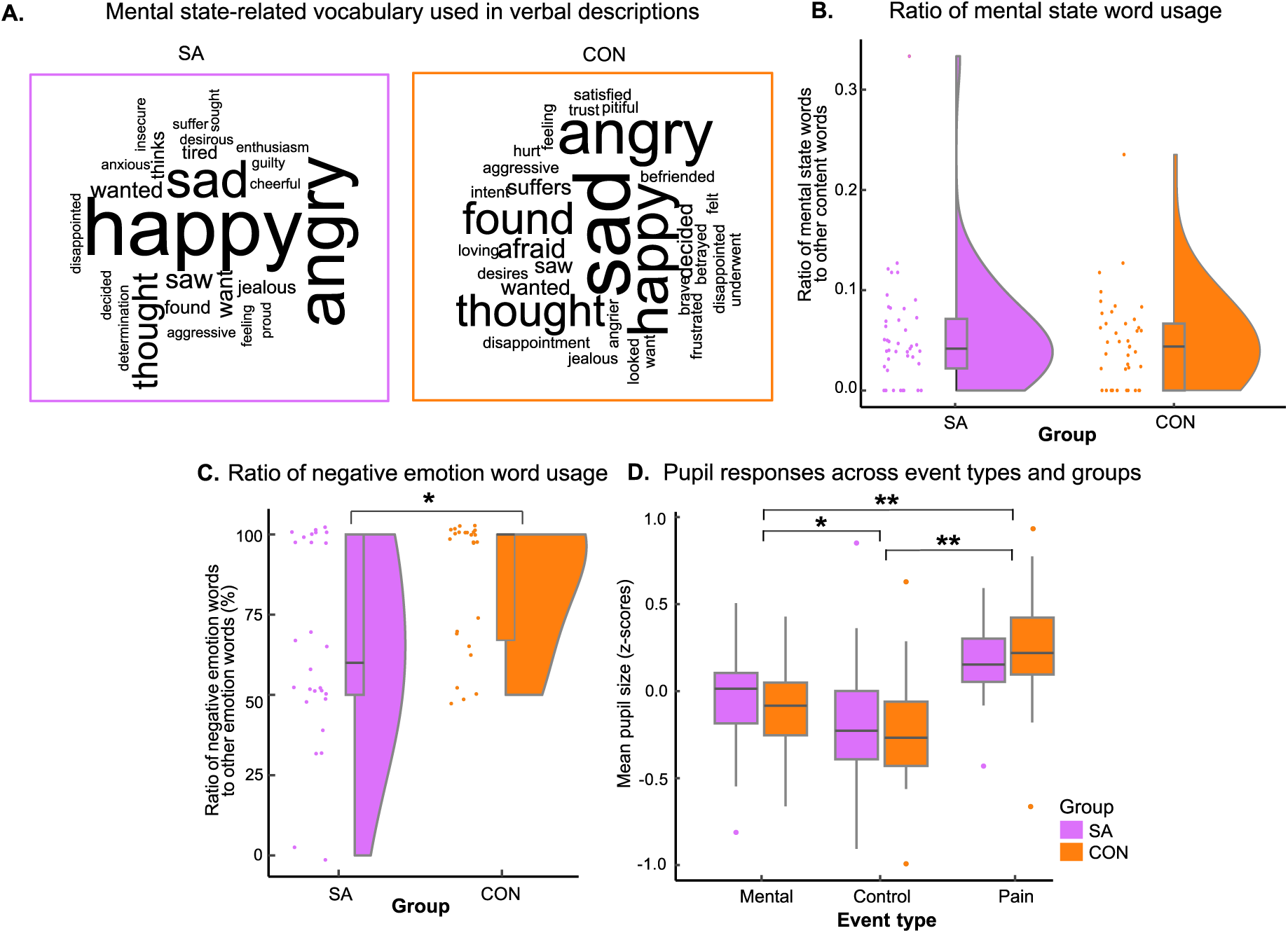
Movie plot descriptions and event-related pupil responses. **A** Word Clouds illustrate the frequency of mental state-related words used by socially anxious and control participants in their movie descriptions. The size of each word is scaled according to its frequency. **B** Proportion of mental state-related words relative to other content words, with no significant differences observed between groups. **C** Proportion of words expressing negative emotions compared to other emotion words, showing a lower incidence of negatively charged words among socially anxious participants relative to controls. **D** Pupil responses across all event types and participants groups, with significant differences between event types marked by asterisks (\*\**p* < .001, \**p* < .01) and no significant differences between groups. Outliers are represented by dots, while whiskers extend to a 1.5 inter-quartile range.

### Heart rate responses

Heart rate, an indicator of physiological arousal [29], exhibited significant fluctuations throughout the movie (Fig. S1a). However, these changes followed a consistent pattern between socially anxious participants and controls, with no significant differences in heart rate at any point during the film or in average heart rates between the groups (M_SA_= 65.4, M_CON_ = 64.0, *t*(72) = 0.62, *p* = .54). Further analysis of intersubject variability in heart rate also showed no significant differences at any point during the film or in average intersubject correlations between the groups (M_SA_ = 0.20, M_CON_ = 0.20, *t*(72) = 0.10, *p* = .92; Fig. S1b). These findings suggest that physiological arousal levels, as measured by heart rate, were similarly experienced by participants from both groups throughout the viewing.

### Event-related pupil responses

Pupil size, an indicator of cognitive effort [38, 49], varied significantly across the three main event types in the film (*F*(2,152) = 43.8, *p* < .001, BF_Null_ = 0.00; Fig. 2d). *Pain* events elicited the largest pupil dilations (M = 0.21, *z*-score), followed by smaller dilations during *Mental* (M = –0.07) and *Control* events (M = –0.21). However, comparisons of pupil responses between socially anxious and control participants revealed no significant differences (*F*(1,73) = 0.04, *p* = .84, BF_Null_ = 6.75). Additionally, there was no significant interaction between group and event type concerning pupil size (*F*(2,152) = 0.63, *p* = .53, BF_Null_ = 7.30), indicating that both groups exhibited comparable levels of cognitive effort in response to the film’s content.

### Event-related brain responses

Whole-brain analyses revealed significant activation within brain regions comprising the Theory of Mind (ToM) network during *Mental* events compared to *Pain* and *Control* events (Fig. 3a). Prominent areas of activation included the right temporoparietal junction (rTPJ: xyz_MNI_ = [44, –58, 30], both *p*_FWE_ < .001), left temporoparietal junction (xyz_MNI_ = [–44, –60, 28], both *p*_FWE_ < .001), precuneus (xyz_MNI_ = [6, –64, 42], both *p*_FWE_ < .001), and medial prefrontal cortex (mPFC: xyz_MNI_ = [6, 54, 26], both *p_FWE_* < .001). Notably, within this network, the left posterior superior temporal sulcus (pSTS) showed a significant reduction in activation for socially anxious participants compared to controls across both contrasts (Fig. 3b; *Mental* > *Control*: xyz_MNI_ = [–44, –62, 18], *t* = 4.41, *p_FWE_* = .003; *Mental* > *Pain*: xyz_MNI_ = [–46, –62, 18], *t* = 4.00, *p_FWE_* = .009).

**Fig. 3.**
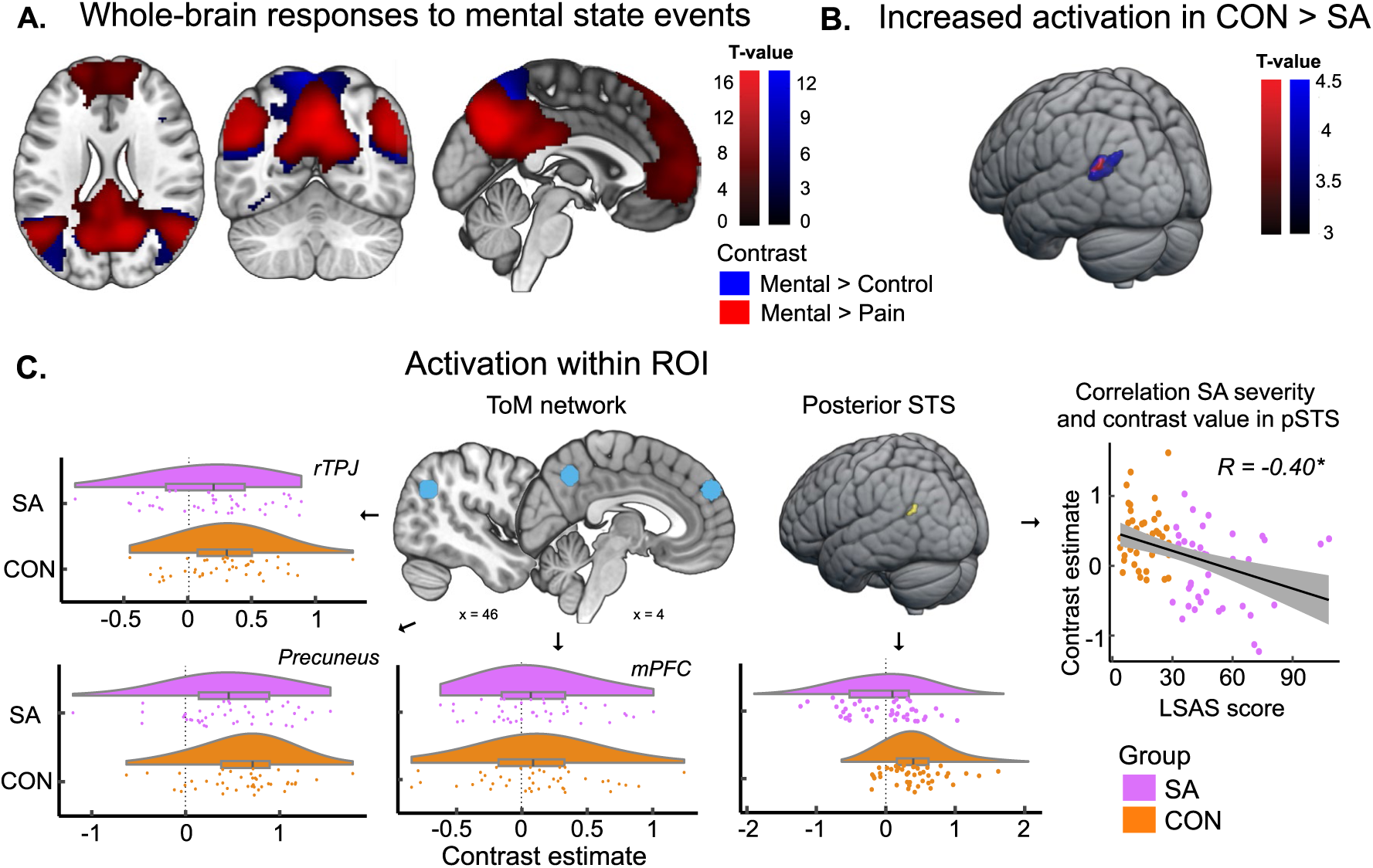
Whole-brain responses to mental state events. **A** Brain regions with significant activation during Mental events as compared to Control and Pain events. **B** Control participants showed greater activation in the left posterior superior temporal sulcus (pSTS) than socially anxious participants across both ToM contrasts. **C** Region of Interest (ROI) analyses underscore the distinct role of the left pSTS within the ToM network in differentiating socially anxious participants from controls. This distinction is supported by a significant correlation between reduced activation in this region and higher levels of social anxiety, as measured by the LSAS.

Region of interest (ROI) analyses reinforced these findings (Fig. 3c). Unlike other regions within the ToM network, the left pSTS displayed clear activation differences between groups, with a Bayes Factor strongly favoring these differences (BF_Group_ > 1085.10). In contrast, primary ToM regions such as the rTPJ, precuneus, and mPFC showed no significant group differences, with Bayes Factors supporting the null hypothesis (rTPJ: BF_Null_ > 2.11; precuneus: BF_Null_ > 1.98; mPFC: BF_Null_ > 4.37). An additional linear regression analysis across all participants linked reduced activation in the left pSTS to higher anxiety levels as measured by the LSAS (r = –0.40, *p* < .001; r_SA_ = –0.11, *p* = .47; r_CON_ = 0.00, *p* = .98), highlighting its unique role within the ToM network in distinguishing socially anxious participants from controls during scenes that involve mental state analysis.

### Movie-driven variability in pupil responses

After conducting event-related comparisons for specific scenes, we examined intersubject variability in responses across the entire film. Dynamic intersubject correlation analysis of the pupil timeseries showed substantial fluctuations during the viewing. For instance, correlations for both groups diminished between 20 and 80 seconds (Fig. S2), during scenes featuring storks flying and clouds transforming into baby animals. While socially anxious participants exhibited lower correlations during these and other intervals of the film, these differences did not reach statistical significance at any point during the viewing.

### Movie-driven variability in brain responses

Using dynamic intersubject correlation analysis on the fMRI timeseries, we identified three spatiotemporal clusters that showed differences in brain response variability between socially anxious and control participants (Fig. 4). Socially anxious participants demonstrated heightened variability in two clusters. Cluster 1 spanned the initial two-thirds of the film (*cluster stat* = 8585, *p* = .003) and included peaks in the right superior parietal lobule (xyz_MNI_ = [24, –52, 74], *t*_max_ = 5.54, *t*_sum_ = 105.26) and both the left and right middle temporal gyrus (xyz_MNI_ = [–60, –10, –22], *t*_max_ = 4.06, *t*_sum_ = 94.37; xyz_MNI_ = [67, –40, 10], *t*_max_ = 2.15, *t*_sum_ = 95.64). Cluster 2 emerged in the final third of the film (*cluster stat* = 4497, *p* = .02), with notable activity in the left supplementary motor area (xyz_MNI_ = [0, 8, 62], *t*_max_ = 4.39, *t*_sum_ = 56.54), medial prefrontal cortex (xyz_MNI_ = [–6, 50, 8], *t*_max_ = 6.77, *t*_sum_ = 56.01), and precentral gyrus (xyz_MNI_ = [–54, 2, 20], *t*_max_ = 3.84, *t*_sum_ = 51.21). Conversely, a third cluster (Cluster 3) exhibited reduced variability in socially anxious participants throughout the film (*cluster stat* = –8478, *p* = .002), with peaks in the right middle occipital lobe (xyz_MNI_ = [42, –70, 8], *t*_min_ = – 3.96, *t*_sum_ = 110.16), inferior frontal gyrus (xyz_MNI_ = [30, 32, –10], *t*_min_ = –4.91, *t*_sum_ = 108.48), and superior temporal gyrus (xyz_MNI_ = [48, 2, –10], *t*_min_ = –3.81, *t*_sum_ = 97.43).

**Fig. 4.**
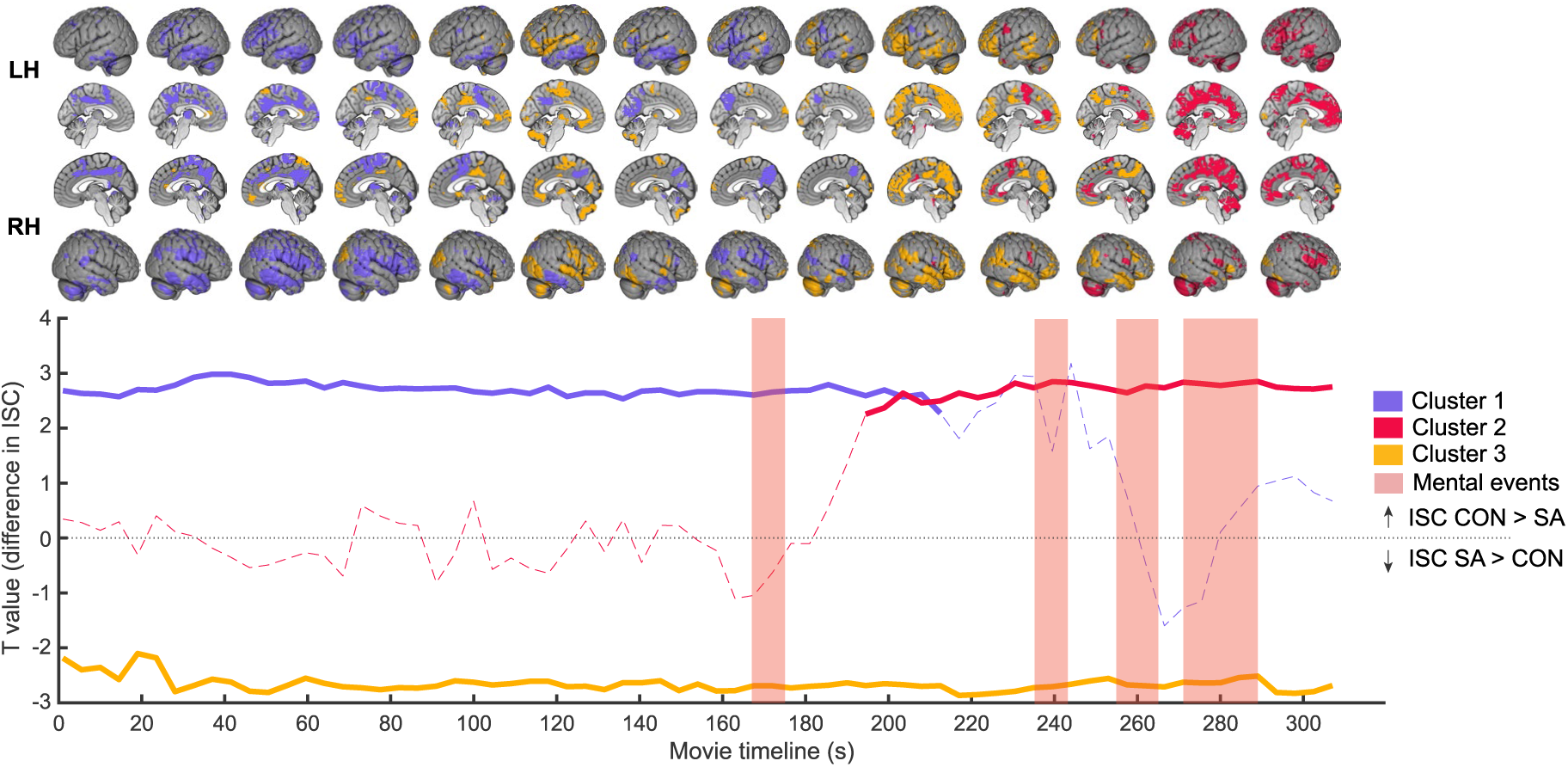
Movie-driven variability in brain responses. Dynamic intersubject correlation analysis identified three spatiotemporal brain clusters that exhibited differences in brain response variability between socially anxious and control participants. In Clusters 1 and 2, socially anxious participants exhibited lower correlations, indicative of heightened variability, across extended intervals of the film. Conversely, Cluster 3 showed consistently reduced variability among socially anxious participants throughout the film. Solid lines depict moments of significant variability within the clusters, while dashed lines represent moments of non-significant variability.

These clusters showed modest spatial overlap with the ToM network, with none exceeding 20% of their respective cluster sizes (Fig. 5). To investigate whether variations in brain response variability correlated with differential activation in the left pSTS during mental state scenes, we analyzed the relationships between intersubject correlations in each cluster and participants’ ToM contrast estimates within the social anxiety group. However, none of these correlations achieved statistical significance (all *p* > .20), suggesting no direct relationship between ToM-related activity in the left pSTS and the variability observed in brain responses across other networks.

**Fig. 5.**
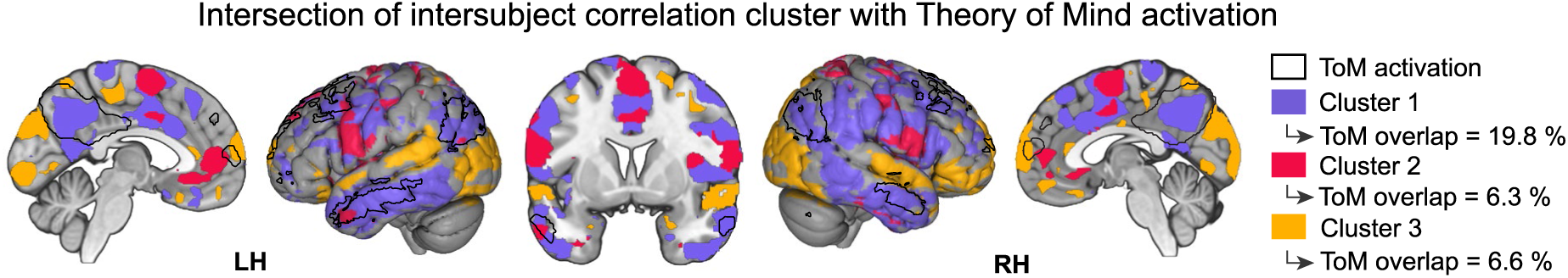
Intersection of neural variability and ToM activation. Overlay of the three brain correlation clusters with activation maps from scenes depicting mental state scenes. Each cluster showed modest spatial overlap with the ToM network, with none of the overlaps exceeding 30% of their respective cluster sizes. For enhanced clarity, clusters are visualized using a cumulative *t*-value threshold of 20 or higher.

### Neural variability in social anxiety versus autism

Given the anatomical similarity between Cluster 1, which exhibited heightened response variability among socially anxious participants, and a previously identified cluster with reduced variability in individuals with Autism Spectrum Condition (ASC) during the same film [28], we conducted a comparative analysis between these groups. The groups were closely matched in terms of gender distribution (62.8% vs. 55.8% women, *p* = .63), age (M_SA_ = 26.3, M_ASC_ = 27.7, *p* = .27), verbal IQ (M_SA_ = 124, M_ASC_ = 126, *p* = .51) and nonverbal IQ (M_SA_ = 102, M_ASC_ = 103, *p* = .79). Levels of social anxiety were also closely matched between the groups (M_SA_ = 51.4, M_ASC_ = 52.4, *p* = 0.84). Our comparative analysis revealed a substantial 60% overlap in brain regions, including the right supramarginal gyrus, right inferior temporal gyrus, anterior midcingulate cortex, and the precuneus (Fig. 6a). Notably, the temporal dynamics within these overlapping regions differed markedly between the groups during the initial two-thirds of the film (Fig. 6b). This divergence suggests that, relative to controls, individuals with social anxiety and those with autism may represent opposite ends of a continuum of neural response variability within this brain network.

**Fig. 6.**
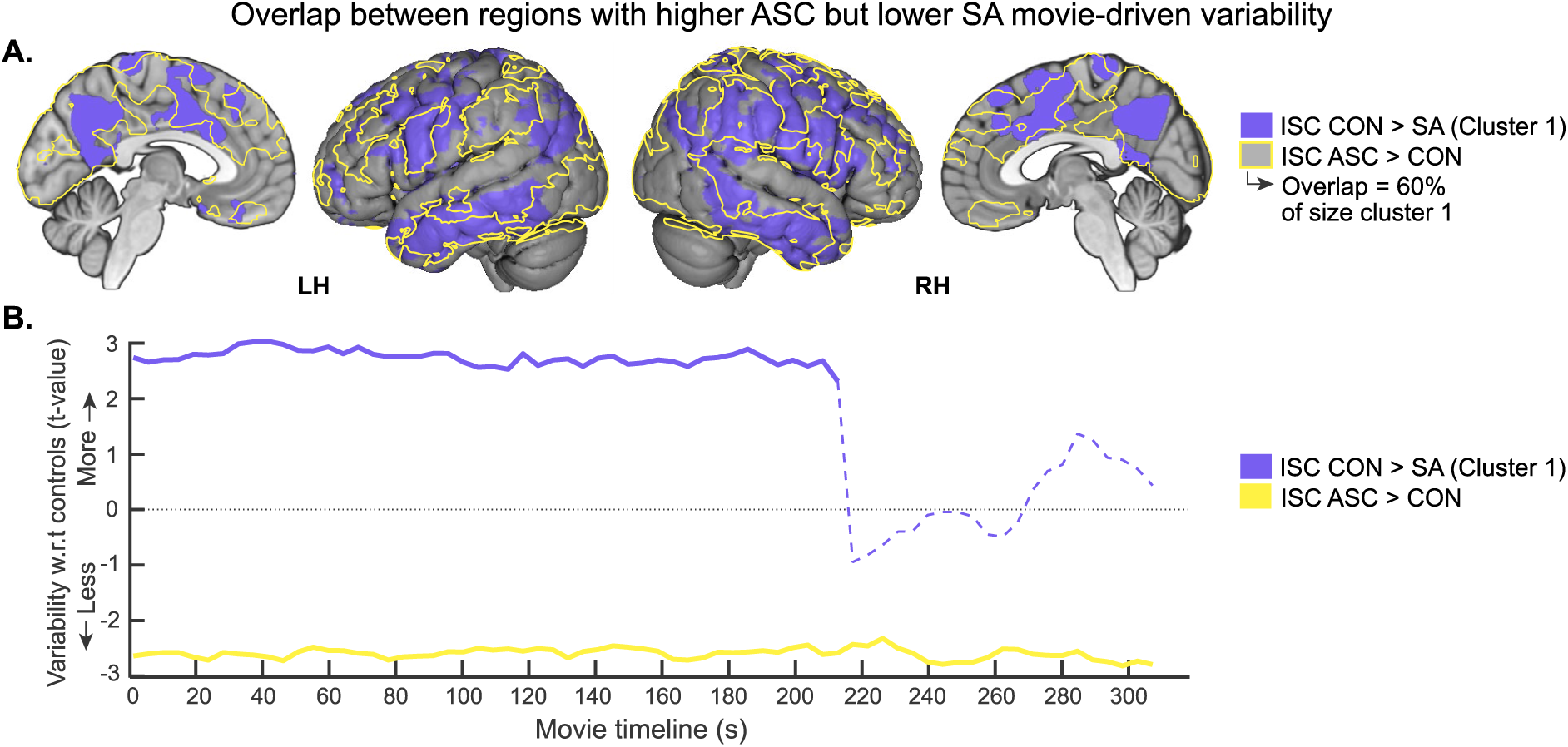
Neural variability in social anxiety versus autism. **A** Overlay of brain correlation Cluster 1, showing heightened variability in socially anxious individuals, with a previously identified cluster that exhibits reduced variability in autistic individuals during the same film. The overlap constitutes 60% of the total cluster size. **B** Temporal dynamics within the overlapping voxels reveal the most significant divergence during the initial two-thirds of the film.

## Discussion

By employing an animated film known for its rich socioemotional dynamics and ability to engage the ToM network during specific scenes, this study provides neural evidence that supports and refines existing cognitive-behavioral models of social anxiety. While overall ToM network activity was similar across groups, socially anxious participants showed significantly reduced activation in the left posterior superior temporal sulcus (pSTS) compared to controls matched for gender, age, and intelligence. Notably, there was an inverse relationship between activity in the left pSTS and participants’ levels of social anxiety, supporting theories that highlight ToM as a fundamental element in SAD [4, 8]. Additionally, intersubject correlation analysis identified a distinctive brain response pattern in regions not directly associated with ToM during film viewing. This pattern, which emerged without corresponding changes in heart rate or pupil responses, indicates a neural processing bias that persists even in non-evaluative settings. Together, these results establish a neural basis for ToM alterations and broader interpretive biases in social anxiety, suggesting important new targets for intervention [14, 15].

Our findings add a neural dimension to the existing behavioral evidence that individuals with SAD underperform on tests of emotion and mental state recognition [8]. For participants with low social anxiety, the left pSTS showed spontaneous activation during scenes depicting characters’ thoughts and emotions. In sharp contrast, participants with high social anxiety did not demonstrate such increased responsiveness in the left pSTS during these scenes, even though activation levels across the broader ToM network remain similar. Furthermore, these ToM alterations manifested in an environment devoid of explicit task demands or overt evaluative pressures, as evidenced by matched levels of physiological arousal and cognitive effort across both high and low anxiety groups. This discrepancy in ToM processing within the left pSTS under non-evaluative conditions suggests that ToM alterations are an inherent feature of social anxiety, not merely a byproduct of evaluative pressures or fear symptoms [4].

The pSTS serves as a critical node within a social cognitive hub centered around the temporoparietal junction (TPJ), which integrates sensory inputs from the external environment with internally generated models of social situations [50]. With functional connections to nearby sensory systems, including motion-sensitive area MT and the fusiform face area (FFA), the pSTS is well-equipped to interpret complex dynamic social scenes involving speech, facial expressions, and other pertinent social stimuli [51–56]. The left pSTS, in particular, is believed to play a crucial role in recognizing others’ emotional states [57, 58]. Consequently, the reduced activation in this region observed among individuals with high social anxiety may reflect a diminished tendency to integrate the emotional content presented in the film with their internal social narratives [59]. This hypothesis is partially supported by post-viewing assessments, where participants with high social anxiety used fewer negative emotion words to describe the film’s plot compared to those in the low anxiety group, suggesting a potential shift in how social events are integrated within narratives.

Consistent with earlier neuroimaging studies of children [24], our results demonstrate both heightened and diminished neural response variability in individuals with high social anxiety during film viewing. Sensory regions such as the bilateral visual and auditory cortices (Cluster 3) showed reduced variability, indicating a more uniform response within these areas among socially anxious individuals. Conversely, two clusters displayed heightened variability, each marked by distinct spatiotemporal patterns in higher-order cortical regions. The first of these clusters became prominent during the climactic final third of the film and included the medial prefrontal cortex (Cluster 2), a region previously identified for its distinct responses in SAD in social task settings [13, 60]. The second cluster was evident during the initial two-thirds of the film and involved the superior parietal lobe and middle temporal gyrus bilaterally (Cluster 1), areas crucial for interpreting socioemotional behaviors [61–64]. These patterns emerged without corresponding changes in pupil size, suggesting an inherent neural processing bias in social anxiety, rather than one driven by conscious effort.

The anatomical extent of Cluster 1 shows considerable overlap with a network demonstrating reduced variability in viewers with autism watching the same film [28]. Detailed analysis revealed a 60% overlap in key areas involved in narrative comprehension and plot development, including the right supramarginal gyrus, inferior temporal gyrus, anterior midcingulate cortex, and the precuneus [65, 66]. The most notable divergence in neural variability within these regions occurred during the initial two-thirds of the film (see Fig. 6), a crucial period for narrative development that likely varies among viewers. This observation suggests that individuals with social anxiety, along with controls and those with autism, may represent a continuum of cognitive engagement with socioemotional narratives. Consistent with this, studies have shown that children with autism are less likely to weave causal explanations of internal states into their personal narratives [67, 68]. For instance, autistic children may use fewer phrases that elucidate the cause or motivation behind events, such as “the boy was mad because the dog broke the jar.” By extension, individuals with social anxiety may demonstrate heightened sensitivity to the broader narrative implications of social events.

The divergence in neural variability observed cannot be exclusively attributed to differences in social anxiety levels, as both the autism and social anxiety groups had comparably high LSAS scores, while the control group scored lower. Despite similar LSAS levels, there were no consistent patterns in neural response variability among these groups compared to the control group. This suggests that the underlying mechanisms of social anxiety may vary across these two groups. Relatedly, social anxiety in autism tends to be more prevalent among older, high-functioning individuals, likely due to increased awareness of social challenges [30]. In contrast, in SAD, the anxiety is likely tied to a shift from basic sensory processing to complex higher-order cognitive functions, consistent with an intrinsic interpretive bias. This distinction is corroborated by eye-tracking studies that assess responses to social stimuli, such as human eyes, in individuals with autism and social anxiety [69, 70]. In these studies, autistic traits correlate with delayed orientation toward the eyes, indicative of a diminished initial salience of human eyes. Meanwhile, traits of social anxiety affect how quickly individuals avert their gaze from the eyes once engaged, reflecting an anxiety-driven avoidance in later processing stages. Collectively, these insights paint a multifaceted picture of social anxiety, illustrating that its underlying mechanisms cannot be unambiguously captured across different diagnoses through self-report measures like the LSAS. Nonetheless, distinct neural patterns are evident and offer valuable new avenues for developing targeted interventions.

A notable finding from this study was that individuals with high social anxiety used fewer negative emotion words in their plot descriptions, diverging from the commonly observed negativity bias in SAD [19]. This occurred even as the total number of mental and emotion-related words used matched that of the low social anxiety group. Several factors might account for this unexpected pattern. First, the offline nature of the assessment and the heightened sensitivity of socially anxious individuals to evaluative pressure might have compelled them to project a more positive self-image. As a safety behavior, a strategy often used to avoid negative evaluations, these individuals may have steered clear of mentioning negative events or emotions depicted in the film [71, 72]. Second, the film’s positive resolution might have overshadowed earlier negative emotions, influencing more positive recall in their plot descriptions. Third, the typical negativity bias associated with social anxiety may be more pronounced in contexts related to self-evaluation [23, 73–75], rather than in interpreting the social performances of others or fictional characters. Future research that combines real-time ratings with neural measures could help disentangle these possibilities.

Lastly, our findings revealed no direct relationship between brain response variability and ToM activation concerning the left pSTS. Although these phenomena involve distinct brain networks, a connection might have been conceivable within the framework of cognitive-behavioral models [14, 15]. For instance, an increased allocation of attentional resources toward monitoring the social narrative, reflected in heightened neural variability, could potentially compete with the resources necessary for interpreting social stimuli, potentially manifesting as reduced activity in the left pSTS. While our study did not detect such a link, future work that manipulates performance monitoring and social task demands could shed light on this hypothetical trade-off between interpretive biases and ToM engagement.

In conclusion, our study demonstrates that socially anxious individuals exhibit reduced spontaneous activation in the left pSTS when viewing movie scenes that depict mental states. This reduction adds a neural dimension to behavioral evidence that these individuals underperform in tests of emotion and mental state recognition, while further illustrating that this ToM alteration persists even in non-evaluative settings. Additionally, our findings delineate a distinct neurocognitive signature characterized by shifts in neural variability, with sensory areas displaying more uniform responses and higher-order cortical regions showing heightened intersubject variability during movie viewing. This signature differentiates individuals with social anxiety from controls, underscoring the importance of incorporating dynamic narratives into research on social perception. By the same token, these results endorse the use of movie-based paradigms to advance our understanding of social anxiety and to pinpoint new targets for intervention. It remains to be seen whether this neurocognitive signature extends to other sociopsychiatric conditions beyond autism, potentially broadening the scope and impact of this research approach.

## Acknowledgements

This work was supported by the NWO Gravitation Grant (024.001.006) to the Language in Interaction Consortium.

## Supplementary Information

**Table S1.** Results of the within-subject fMRI analyses related to the main effects of event type (Mental, Pain, Control).

| *Contrast* | *Anatomical location of peak voxel* | *Cluster size* | *MNI coordinates* |  |  | *T-value*<br>(df = 1, 84) |
| --- | --- | --- | --- | --- | --- | --- |
|  |  |  | *x* | *y* | *z* |  |
| Mental > Control | Right precuneus | 17625 | 2 | -54 | 46 | 13.41 |
|  | Left fusiform gyrus | 2207 | -28 | -48 | -8 | 10.07 |
|  | Right middle temporal gyrus | 1897 | 56 | 2 | -26 | 9.57 |
|  | Left middle temporal gyrus | 1862 | -52 | 4 | -28 | 8.81 |
|  | Left middle frontal gyrus | 5636 | -32 | 10 | 48 | 6.37 |
|  | Right postcentral gyrus | 396 | 42 | -22 | 38 | 5.40 |
|  | Left middle orbital gyrus | 240 | 2 | 62 | -10 | 4.80 |
|  | Right thalamus | 305 | 18 | -26 | 10 | 4.68 |
| Mental > Pain | Right precuneus | 36624 | 6 | -64 | 42 | 16.68 |
|  | Left middle temporal gyrus | 2211 | -54 | 8 | -32 | 10.74 |
|  | Right middle temporal gyrus | 1425 | 60 | -10 | -20 | 8.36 |
|  | Right parahippocampal gyrus | 304 | 20 | -36 | -12 | 5.94 |

**Fig. S1.**
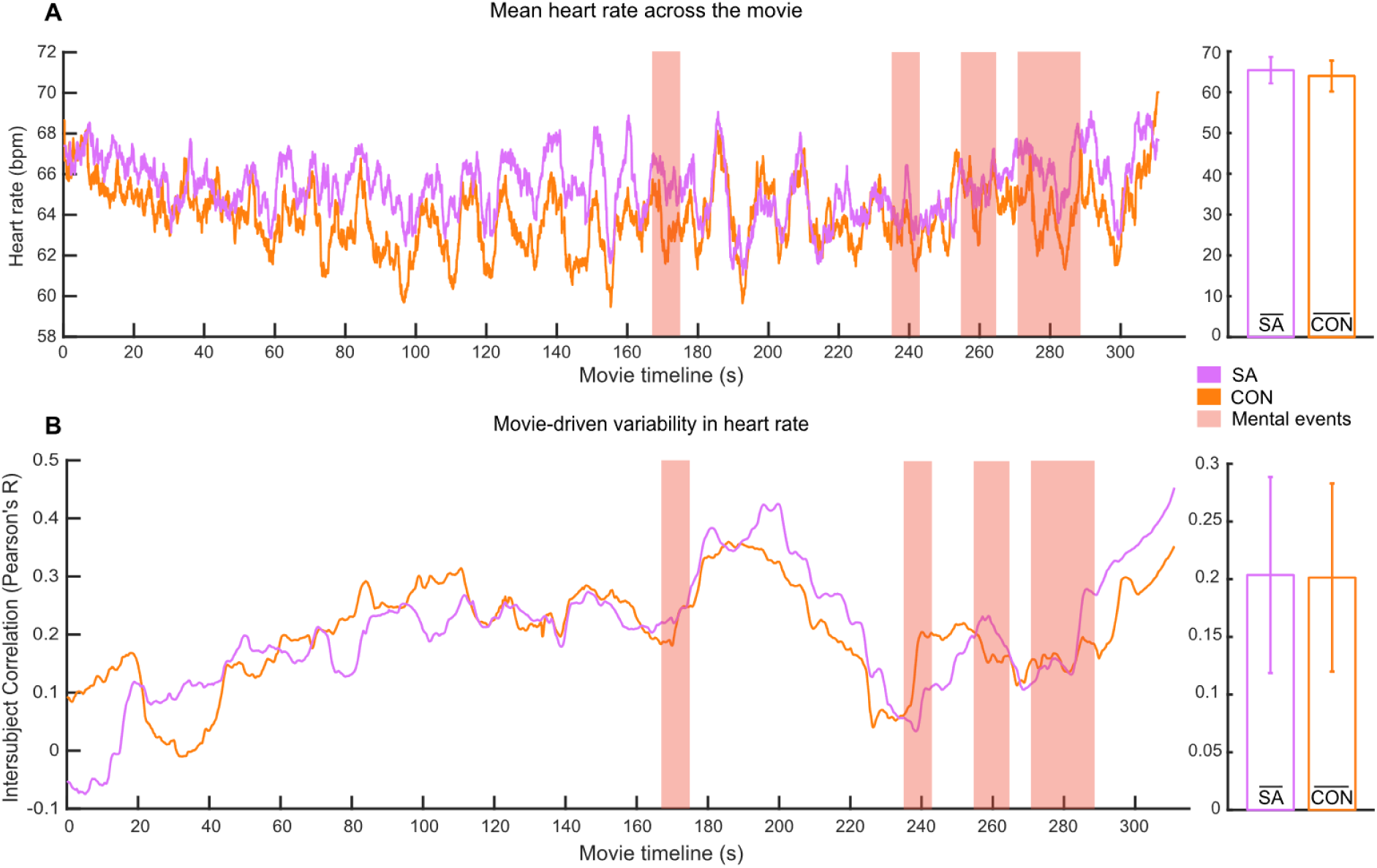
Heart rate metrics and intersubject variability during movie viewing. **A** Mean heart rate measured across the duration of the movie for both socially anxious and control participants. Adjacent histograms show the aggregated mean heart rate throughout the entire film for each group, with error bars representing 95% confidence intervals. **B** Average intersubject correlation coefficients illustrate the intersubject variability in heart rate response during the movie among socially anxious and control participants. There were no significant differences detected in either metric between the two groups.

**Fig. S2.**
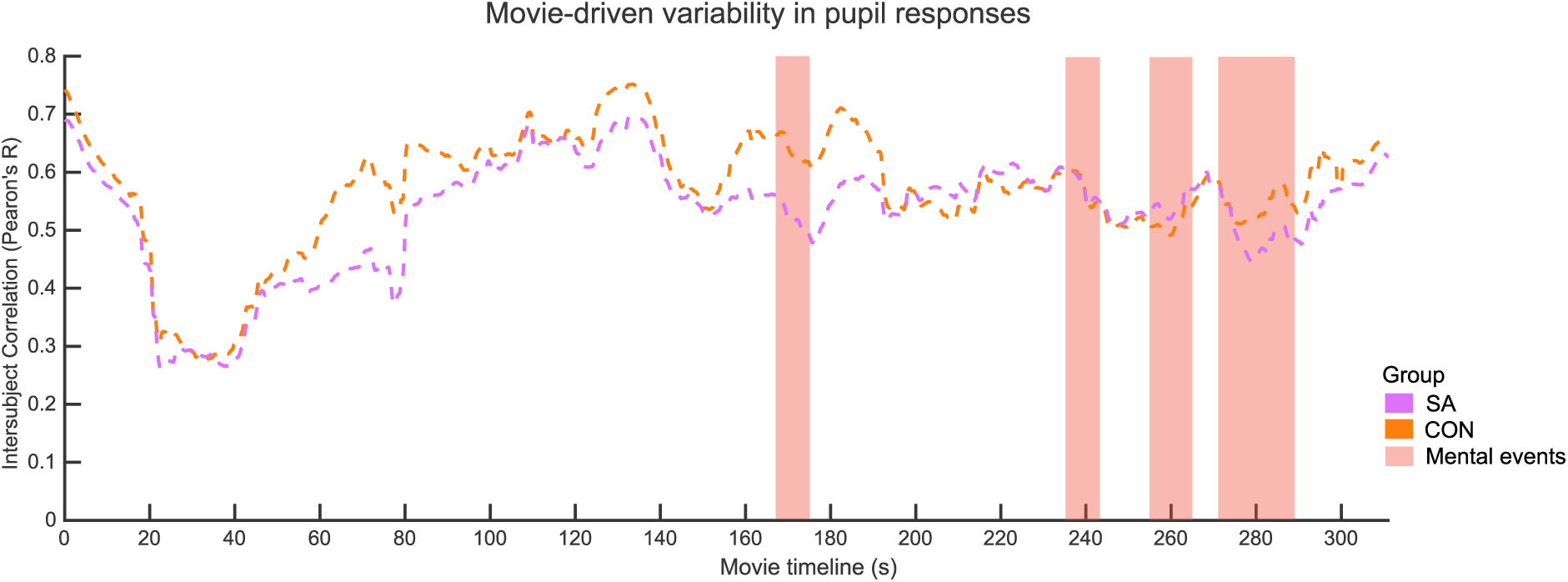
Movie-driven variability in pupil responses. Dynamic intersubject correlation analysis revealed substantial fluctuations in variability throughout the film. Socially anxious participants exhibited lower correlations, indicative of higher variability in pupil responses compared to controls. Nonetheless, these differences did not reach statistical significance at any point during the viewing.

